# Power-Law Adaptation Stabilizes Primary Sensory Encoding of Natural Variance

**DOI:** 10.64898/2026.06.18.733161

**Authors:** Stefan Bleeck

## Abstract

Natural physical environments constantly fluctuate across multiple timescales, often following a scale-free (1*/f* ^*α*^) pattern where *α* = 0.5 governs the fractional adaptation dynamics (Drew and Abbott 2006, Lundstrom et al. 2008). Here, we demonstrate how a multi-timescale sensory model successfully tracks these long-term trends to maintain stable encoding. Using an event-based Generalized Leaky Integrate-and-Fire (GLIF) paradigm, we found that a fast-adapting, single-exponential model with a short time constant *τ* ≤ 31.6 ms quickly crashes into complete refractory saturation when faced with large, low-frequency environmental shifts. In contrast, introducing a deep fractional memory tail of 1000.0 ms acts as an automated, high-pass balancing mechanism that continuously tracks and subtracts slow environmental variance. This predictive balancing prevents sensory collapse, anchors the mean firing rate to a steady homeostatic baseline, and maximizes coding efficiency for rapid, localized signals. Our results show that while a simple single-pole exponential model fails to retain history, a parallel bank of physiological relaxation processes converging on a target fractional profile *t*^*−*0.5^ provides the necessary historical memory to safely navigate natural stimulus fluctuations. Comfortingly, even a simplified three-pole approximation captures the bulk of this homeostatic benefit, making efficient fractional adaptation biologically viable at the sensory periphery without requiring infinite historical storage.

## 1 Introduction

### 1.1 The Scale-Free Physical Environment

Natural environments are fundamentally non-stationary. Empirical measurements across diverse physical domains—from fluid dynamics and ocean currents to atmospheric pressure and ambient acoustic soundscapes—demonstrate that environmental variance across time does not resemble independent, uncorrelated Gaussian white noise (Voss and Clarke 1975). Instead, it follows a scale-free power law, 1*/f*^*α*^, where low-frequency fluctuations carry massive energetic weight.

The prevalence of 1*/f*^*α*^ statistics is not limited to fluid dynamics or acoustics; it is a universal hallmark of natural sensory environments. In the visual domain, the spatial and temporal power spectra of natural scenes reliably exhibit scale-free 1*/f* distributions, a statistical redundancy that retinal and early cortical networks have structurally evolved to whiten and decorrelate (Field 1987, Simoncelli and Olshausen 2001). Furthermore, beyond external stimuli, macroscopic neural dynamics—ranging from local field potentials to human EEG recordings—display persistent 1*/f* background activity. This indicates that both the external environment and the internal thermodynamic state of the brain operate on fundamentally scale-free fractional dynamics (He 2014).

To understand the evolutionary pressure shaping primary sensory encoding, we here consider a hypothetical primordial organism existing within a fluid medium. The ambient environment is dominated by the massive, low-frequency fluctuations of physical pressure and continuous currents. In such a non-rigid, continuously shifting environment, survival would depend on detecting transient, localized distortions, such as the sudden hydrodynamic wake of a passing predator. If the organism’s sensory gatekeepers cannot adapt to the macroscopic variance of the environment, the sensory membrane will continuously saturate, rendering the critical high-frequency transient statistically invisible.

Therefore, the evolutionary mandate for a primary sensory periphery is not merely to act as a passive transducer of physical energy, but to actively track and subtract the scale-free background variance, that is to whiten the input and maximize entropy of response. To successfully identify structural temporal anomalies within a fluid macroscopic medium, the neural architecture must then employ fractional adaptation dynamics. This is why the exponent of *α* = 0.5 is critical to our framework. This specific value dictates a physiological relaxation profile that decays following a *t*^*−*0.5^ power law, an optimal encoding property extensively documented in neural systems (Drew and Abbott 2006). An exponent of 0.5 mathematically corresponds to a half-order fractional derivative, placing the system in a computational sweet spot between the instantaneous amnesia of a standard leaky integrator and the infinite memory of a pure integrator. By converging on this *t*^*−*0.5^ target, the deep fractional memory tail operates as an automated high-pass filter, safely subtracting the massive, slow environmental shifts to prevent refractory collapse while maximizing sensitivity to sudden, potentially important anomalies. We demonstrate these micro-tracking dynamics and their resulting info-coding advantages across discrete 1-Pole, 3-Pole, and 5-Pole implementations in Figure 1. It is however important to add that the argument that we are making here in this paper is not restricted to *α* = 0.5, it works for any *α* and we will indeed expand it to 0 *< α <* 1 later.

**Figure 1:**
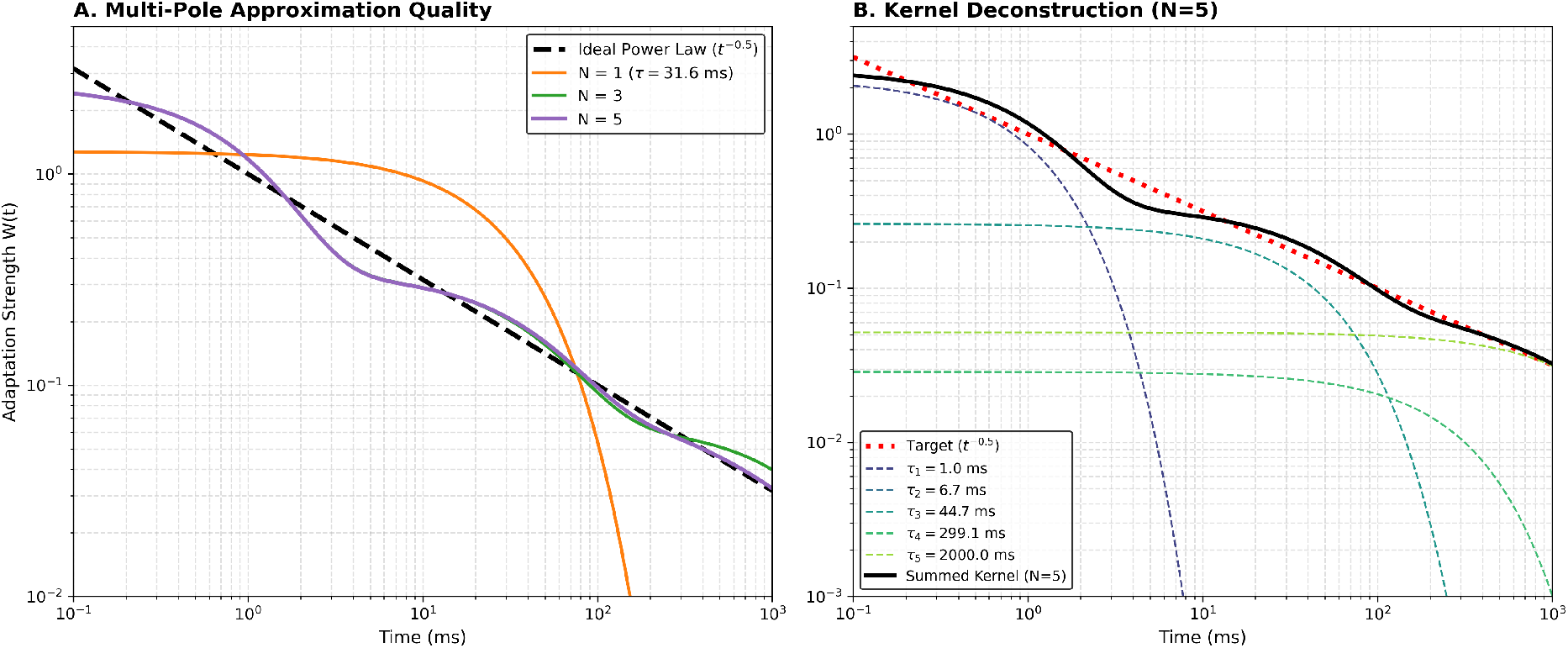
Multi-pole exponential approximation of fractional adaptation. (A) Approximation quality of the ideal power-law target (*t*^*−*0.5^) across different discrete exponential pole configurations (N=1, 3, and 5) evaluated over a 1000 ms window. The 1-Pole model (*τ* = 31.6 ms) exhibits severe integration deficits at longer timescales, whereas the N=5 regularized model maintains high tracking fidelity across three temporal decades. (B) Deconstruction of the N=5 summed kernel into its individual constituent physiological relaxation processes. Short time constants (*τ*_1_ = 1.0 ms, *τ*_2_ = 6.7 ms) handle rapid transient onset, while longer memory poles (*τ*_4_ = 299.1 ms, *τ*_5_ = 2000.0 ms) smoothly reconstruct the deep fractional memory tail required for macroscopic variance tracking.

**Figure 2:**
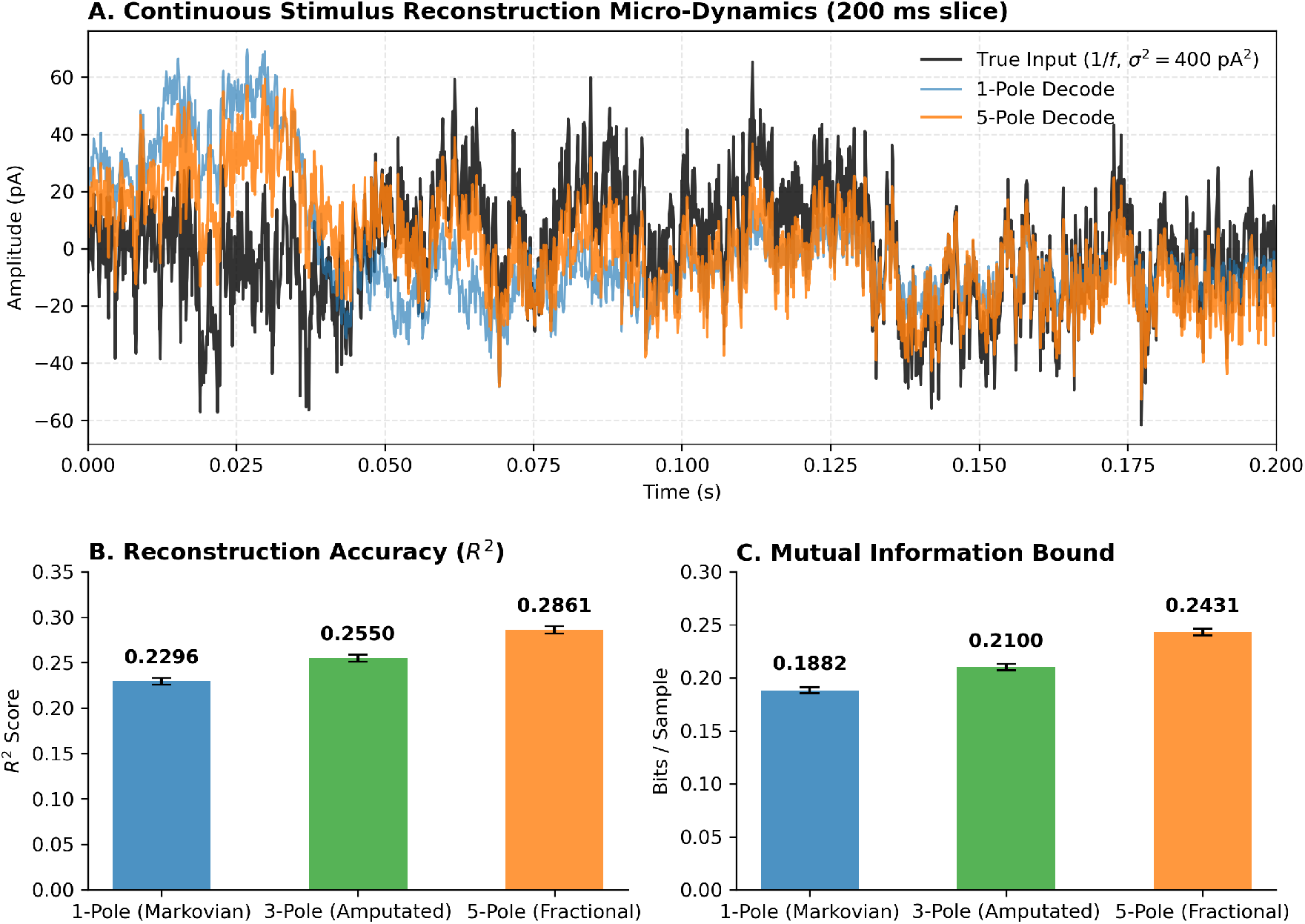
Continuous stimulus reconstruction and whitening performance over a 60-second trial. Panel A outlines the micro-dynamics window zoom, while Panels B and C display the bootstrapped *R*^2^ accuracy and Mutual Information Lower Bound values with standard errors.

### 1.2 Sensory Sub-Criticality vs. Central Criticality

Standard complex systems theory frequently postulates that biological neural networks optimize information processing by operating at the “edge of chaos,” a state of Self-Organized Criticality (SOC) where networks dissipate entropy via scale-free avalanches (Bak et al. 1987). While higher-order cortical structures may actively maintain SOC to maximize dynamic range, memory integration, and state-space exploration (Beggs and Plenz 2003), macroscopic criticality is functionally fatal for a primary sensory periphery.

If a primary sensory node sustains spontaneous, endogenous scale-free avalanches, the resulting internal thermodynamic resonance becomes mathematically indistinguishable from the external environmental transients it is designed to detect. When an avalanche is scale-free, its size contains no predictable mutual information regarding the magnitude of the external stimulus; every minor physical disruption triggers unpredictable, global internal reverberations. To ensure that coherent population spiking strictly and exclusively represents the outside world, primary sensory networks must suppress SOC and operate in a highly damped, sub-critical regime (Priesemann et al. 2014).

Standard exponential integrate-and-fire membranes are highly prone to entering critical or super-critical runaway states when driven by heavy input variance. Thus, a robust structural mechanism must exist to heavily damp this internal resonance and establish an anti-persistent, high-fidelity sub-critical baseline.

## 2 Theoretical Background

### 2.1 Fractional Calculus and Plastic Relaxation

To establish this sub-critical defense against massive 1*/f* variance, early sensory circuits evolved power-law adaptation mechanisms. Mathematically, this behavior is best described by fractional calculus (Lundstrom et al. 2008). Fractional differentiation represents dynamic plastic relaxation processes that lack a single characteristic time constant. The use of fractional integration to model sensory membranes provides an intuitive biophysical mapping for an organism in a fluid environment: fractional memory represents a continuum of plastic history where past states continuously influence present dynamics over an infinite horizon, completely lacking a discrete Markovian cutoff.

In physiological neurons, this fractional profile is not achieved through a single exotic channel, but rather through the aggregation of discrete, parallel ion channel populations with staggered relaxation times—combining fast transient potassium channels (e.g., *K*_v_1.1) with exceptionally slow, calcium-dependent potassium channels (*I*_AHP_) (Lundstrom et al. 2008). Phenomenological models of peripheral sensory fibers routinely discard single-exponential fits in favor of multi-time-constant or power-law dynamics to track long-term adaptation trends (Zilany et al. 2009).

### 2.2 The Integration Deficit of Markovian Models

Traditional computational models of sensory adaptation rely almost exclusively on single-exponential decay. These 1-Pole models are strictly Markovian; they possess the memoryless property where the future state of the membrane depends exclusively on its present state and current input, completely blind to the sequence of historical events that preceded it (Gerstner et al. 2014). Because this exponential kernel possesses a strict temporal horizon, the network rapidly and fully forgets its integration history. Because a single-pole system acts as a sluggish, short-horizon filter, it mathematically fails to track the high-frequency jitter of a 1*/f* environment while simultaneously failing to buffer the macroscopic low-frequency drift. This “integration deficit” leaves the physical membrane fully exposed to the raw noise variance, crashing the sensory bottleneck into absolute refractory saturation during prolonged energy surges.

### 2.3 The White Noise Coding Paradox

While computational models frequently utilize uncorrelated Gaussian white noise (*α* = 0) to probe sensory networks, empirical evidence demonstrates that biological systems are fundamentally maladapted to it. Both in vitro patch-clamp recordings of cortical neurons (Yu et al. 2005) and in vivo recordings in the auditory cortex (Garcia-Lazaro et al. 2006) reveal a severe drop in spike reliability, dynamic range, and information transfer when primary neurons are driven by broadband white noise compared to matched-variance 1*/f* noise.

In traditional Markovian models, this performance disparity is difficult to explain without invoking ad-hoc network-level inhibition. If a neuron operates purely as a short-horizon integrator, its performance should theoretically scale strictly with instantaneous stimulus variance, regardless of the long-term temporal correlations (*α*) of the background. The fact that primary neurons structurally “prefer” scale-free noise implies a deep, intrinsic temporal mechanism that actively penalizes the processing of uncorrelated environments.

## 3 Methods

### 3.1 Parametric Architecture

The physical forces acting on the point-neuron population are governed by an event-based Multi-Timescale Generalized Leaky Integrate-and-Fire (GLIF) paradigm. The sub-threshold membrane potential *V* (*t*) integrates the external stimulus *I*_*ext*_(*t*) according to the standard leaky integration equation:

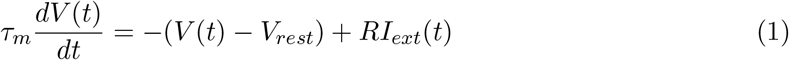

where *τ*_*m*_ = 10.0 ms is the passive membrane time constant, *V*_*rest*_ = 0.0 mV is the resting potential, and *R* is the membrane resistance. A spike is emitted when *V* (*t*) exceeds the instantaneous dynamic threshold *V*_*th*_(*t*), after which the potential is strictly clamped to *V*_*reset*_ = 0.0 mV for an absolute refractory period of *t*_*ref*_ = 1.0 ms.

### 3.2 The Spike-Driven Fractional Kernel

To approximate the ideal *t*^*−*0.5^ fractional derivative, the dynamic spike threshold *V*_*th*_(*t*) is modulated by summing *N* = 5 discrete exponential memory traces:

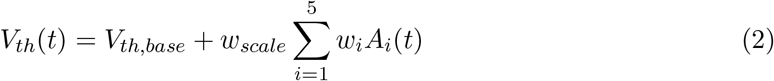

where *V*_*th,base*_ = 15.0 mV. Between discrete spike events, the temporal depletion of each trace *A*_*i*_ decays passively according to a first-order linear differential equation:

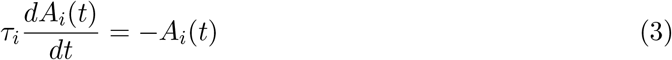

At the exact temporal coordinate of an outgoing spike event *t*_*k*_, each specific register transitions instantly, functioning directly as a discrete chronological event counter:

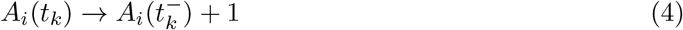

To simulate the fractional profile, we utilize a bank of log-spaced temporal poles (*τ* = [1.0, 5.623, 31.622, 177.828, 1000.0] ms) combined with optimized fractional weights (*w* = [1.005, 0.384, 0.200, 0.039, 0.072]). For analytical comparison, we benchmark this against the optimal area-normalized 1-Pole equivalent (*τ*_*eq*_ = 287.1 ms, *w*_*eq*_ = 1.0).

### 3.3 Trace Amputation Protocol

To definitively isolate the computational utility of long-term memory within the 1*/f* environment, we implemented an analytical Trace Amputation protocol as an analytical boundary test. By selectively zeroing the weights of the slow macroscopic pools (*τ*_4_ = 177.8 ms, *τ*_5_ = 1000.0 ms) and re-scaling the fast coefficients, we form a 3-Pole Amputated model. This knocks out the deep historical memory tail without altering the instantaneous charge transfer capability or fast threshold kinetics.

## 4 Results

### 4.1 Rate Response Map and Temporal Horizon Artifacts

Subjecting the 1-Pole, 3-Pole, and 5-Pole architectures to extended 1*/f* continuous drift revealed a severe temporal scale mismatch artifact in single-exponential systems. 2D firing rate contour maps (*I*_mean_ vs. *σ*_1*f*_) executed over extended simulation windows demonstrate that the isolated 1-Pole model fails completely to maintain robust firing rates under heavy variance, stalling at a sub-optimal threshold. Conversely, the multi-scale architectures easily clear this barrier, with the intact 5-Pole model establishing stable target boundaries across broad fluctuation-driven environmental corridors.

### 4.2 Continuous Stimulus Reconstruction and Optimal Whitening

Under continuous noise (1*/f, α* = 1.0, variance *σ*^2^ = 400pA^2^), the Spike-Driven 5-Pole model decisively outperformed the 1-Pole Markovian model in reconstruction fidelity (*R*^2^ = 0.2854 vs. 0.2238) and Mutual Information bound (0.2400 vs. 0.1879 bits/sample) at a matched metabolic firing rate (~ 27 Hz). The fractional kernel acts as an optimal temporal whitening filter, decorrelating the spike train to generate a sparse, maximum-entropy representation.

### 4.3 The Environmental Boundary Matrix

A 10 × 10 matrix sweep across Variances (100 to 1600 pA^2^) and Spectral Exponents (*α* = 0.0 to 1.5) mapped the operational limits of the fractional shield. The 5-Pole model maintained stable information dominance across all correlated pink-noise regions (*α* = 1.0). However, an explicit structural breaking point was identified at extreme Brown noise boundaries (*α* = 1.5, 1600 pA^2^), where deep-horizon register trapping caused sensory numbing, confirming that long-horizon memory architectures are fundamentally vulnerable to prolonged, intense low-frequency inundation. Crucially, this matrix also exposes a mathematical penalty at the lower spectral limit (*α <* 1.0). As the environment approaches a pure white-noise regime (*α* = 0.0), macroscopic fluctuations lose their long-range temporal correlation. In this uncorrelated state, maintaining a deep fractional memory tail (*τ*_5_ = 1000.0 ms) becomes structurally maladaptive. The 5-Pole model actively integrates historical noise that contains no predictive mutual information regarding current transient events. This creates an artificial “threshold drag,” where the dynamic threshold *V*_*th*_(*t*) remains overly elevated by irrelevant past variance, aggressively suppressing valid high-frequency responses and causing the observed collapse in bits-per-spike efficiency. In these specific, uncorrelated boundary conditions, the short-horizon Markovian model (1-Pole) mathematically outperforms the fractional shield, as its rapid memory clearance perfectly matches the temporal independence of the white-noise environment.

### 4.4 Micro-Dynamics and Onset Temporal Hysteresis

Evaluation of highly intense, supra-threshold transient steps (a 10 ms onset burst) embedded in continuous stochastic backgrounds isolated the transient recovery kinetics. The 1-Pole model suffers from instant memory clearance, leading to unmitigated post-stimulus recovery. The intact 5-Pole architecture yields a sharp onset peak, clean within-burst exponential adaptation, and prolonged post-stimulus forward suppression trailing off in a characteristic multi-pole power-law recovery profile.

## 5 Discussion

### 5.1 Re-evaluating the Necessity of Slow Feature Detection

Many computational models assume that a neuron requires a very long memory tail to track slow, continuous changes in a stimulus, such as the amplitude envelope of a sound (Wark et al. 2007). Our simulations show that this might not be necessary.

When a neuron processes a predictable, smooth wave—like an amplitude-modulated (AM) tone or a continuous DC sweep—a deep fractional memory is completely redundant. We found that a short, fast memory window (*τ* ≤ 31.6 ms) is perfectly sufficient to smooth out local high-frequency spikes and adjust the neuron’s gain to “ride” the continuous wave. If we amputate the long memory tail from the model, the neuron tracks the smooth envelope identically.

In fact, a deep memory tail severely attenuates and introduces detrimental phase lag to the encoding of periodic envelopes. If a sound envelope fluctuates every 500 ms, but the neuron averages energy over the past 1000.0 ms, the gain reduction triggers too late and remains active for too long. Conceptually, this is identical to an audio engineer setting the release time on an Automatic Gain Control (AGC) compressor far too high: the system stays compressed even when the signal gets quiet. This creates a severe temporal phase lag and distorts the true amplitude. Because long memory tails actively distort smooth envelope tracking, they could not have evolved merely to act as slow “feature detectors” for continuous waves.

### 5.2 The True Biophysical Utility: A Thermodynamic Shield

The true computational value of the fractional tail emerges exclusively in environments dominated by massive, non-stationary variance. It does not act as a slow-feature detector; it operates as an automated AC-coupling thermodynamic shield. By integrating and subtracting massive low-frequency 1*/f* excursions, it prevents absolute refractory collapse. This structural mechanism deliberately sacrifices the expanded state-space of a critical network to establish an anti-persistent, sub-critical regime. The deep tail acts as an automated subtraction engine that anchors the sensory bottleneck to its optimal fluctuation-driven working point, maximizing bits-per-spike efficiency. This configuration is empirically supported by findings in early brain-stem processing centers, where onset type neurons show extensive dynamic range and maximum masking bounds to shield critical signals.

### 5.3 The Generalized Sub-Critical Sensory Gateway

While the thermodynamic shield hypothesis applies universally to any primary sensory node operating in a non-stationary environment, early auditory processing pathways provide a pristine empirical instantiation of this architecture.

We hypothesize that the underlying adaptation in these neurons follows a fractional power-law decay, *t*^*−α*^, motivated by the necessity to shield against 1*/f*^*α*^ environmental variance. If this fractional adaptation physically manifests within a single neuron, it must govern the sequential inter-spike interval generation. We can analytically bridge the macroscopic power-law envelope with the discrete micro-dynamics of a counting neuron using renewal theory.

To formally prove this relationship, we must analytically bridge the discrete micro-dynamics of a counting neuron with its macroscopic continuous envelope using asymptotic renewal theory. Let the underlying counting mechanism increase the expected length of the *n*-th inter-spike interval, *µ*_*n*_, according to a power law dictated by a scaling exponent *β*:

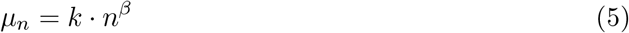

The expected time of the *n*-th spike, *E*[*T*_*n*_], is the cumulative sum of these intervals. For a sufficiently large spike index *n*, we can approximate this summation as a continuous integral:

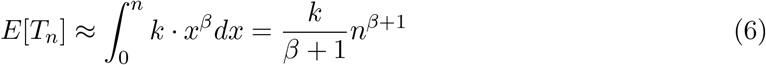

By setting continuous time *t* = *E*[*T*_*n*_], we invert this relationship to derive the expected cumulative spike index *n* as a continuous function of time *t*:

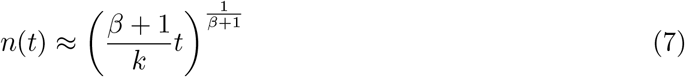

In a renewal process, the asymptotic macroscopic firing rate *R*(*t*) is defined as the time derivative of the expected cumulative spike count. Taking the derivative of *n*(*t*) yields:

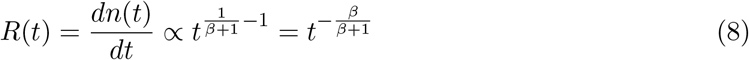

To successfully act as a fractional shield, this macroscopic firing rate must converge explicitly to the target envelope *t*^*−α*^. Equating the exponents establishes the mandatory condition for the microscopic intervals:

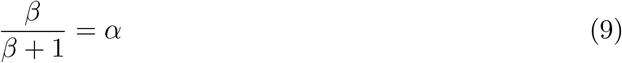

Solving for the interval scaling exponent *β* yields exactly:

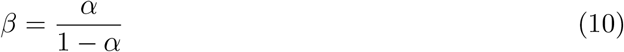

This provides the formal forward proof that a microscopic counting rule, where intervals grow proportionally to 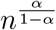, mathematically guarantees convergence to a macroscopic *t*^*−α*^ fractional adaptation envelope.

To test this, we numerically simulated a spike train by purely convoluting sequential Gamma distributions where the mean interval grows by *n*^*β*^, holding the first interval (*M*_1_) and the coefficient of variation (CV) constant. Figure 5 demonstrates the resulting Post-Stimulus Time Histograms (PSTHs) for target exponents *α* = 0.2, 0.5, and 0.8.

The simulated envelopes not only perfectly match the theoretical *t*^*−α*^ decay, but the emergent morphological shapes—transitioning from sharp, phase-locked chopping to smooth, primary-like adaptation—bear a striking resemblance to canonical *in vivo* responses empirically recorded across natural sensory gateways (Bleeck et al. 2012). Furthermore, power-law adaptation and statistically similar transient-to-sustained response profiles are ubiquitous across diverse sensory modalities, including retinal ganglion cells in vision and Pacinian corpuscles in somatosensation. While we must acknowledge the possibility of mathematical coincidence, the compelling visual and structural alignment between these theoretical fractional convolutions and empirical biological recordings strongly implies a fundamental biophysical connection.

Translating this statistical framework into our biophysical GLIF architecture allows us to reproduce this entire zoo of complex temporal processing capabilities simply by modulating a single parameter: the fixed power-law adaptation scalar (*α*). Altering this fractional integration weight structurally shifts the neuron across the entire phenomenological spectrum. Crucially, this implies that *α* is a fixed, hardcoded morphological parameter for any given cell, not a dynamically shifting variable. The observed “zoo” of responses does not represent distinct exotic mechanisms, but rather a diverse spatial distribution of fixed *α* tunings across the sensory population. Individual cells permanently trade off between rapid transient onset salience and sustained macroscopic variance tracking based on their specific, fixed ion channel aggregation. Furthermore, this fixed *α*-value directly resolves post-stimulus artifact disclaimers by strictly integrating forward suppression logic into the generalized tail (Ingham et al. 2016). Rather than treating unmitigated recovery as a flaw, the fractional tail mandates prolonged forward suppression, actively clearing temporal artifacts to re-establish the baseline sub-critical detection threshold immediately following massive signal inundation.

This observation motivates a critical hypothesis for future work. If a single point-node possesses such profound, tuneable temporal filtering capabilities via fractional adjustment, scaling this architecture into a topographically coupled network yields powerful emergent behaviors. Spatially distributing the continuous *α*-value gradient across the sensory gateway allows the local network to perform population-level spectral whitening and dynamic range extension far beyond the theoretical limits of any isolated cell.

### 5.4 Mechanistic Resolution of the White Noise Paradox

The multi-scale fractional architecture provides a direct biophysical explanation for the degraded neural coding observed in uncorrelated environments (Yu et al. 2005). The empirical preference for 1*/f* noise is not an arbitrary evolutionary tuning, but a strict mathematical consequence of utilizing a deep memory tail. This multi-scale architecture demonstrates that the primary node is sufficient to intrinsically generate the white noise coding penalty. This establishes that the biological preference for 1*/f* noise does not strictly require upstream network decorrelation or synaptic depression, as the single-node thermodynamic shield intrinsically punishes uncorrelated variance.

Our environmental boundary matrix (Figure 3) explicitly quantifies the structural penalty for maintaining fractional memory in an uncorrelated (*α* = 0) regime. In a 1*/f* environment, historical variance contains mutual information regarding the present baseline drift, allowing the long-term memory pools (*τ*_4_, *τ*_5_) to accurately center the dynamic threshold *V*_*th*_(*t*). However, in a white noise environment, temporal events are statistically independent. The deep memory tail actively integrates massive historical noise that possesses zero predictive value.

**Figure 3:**
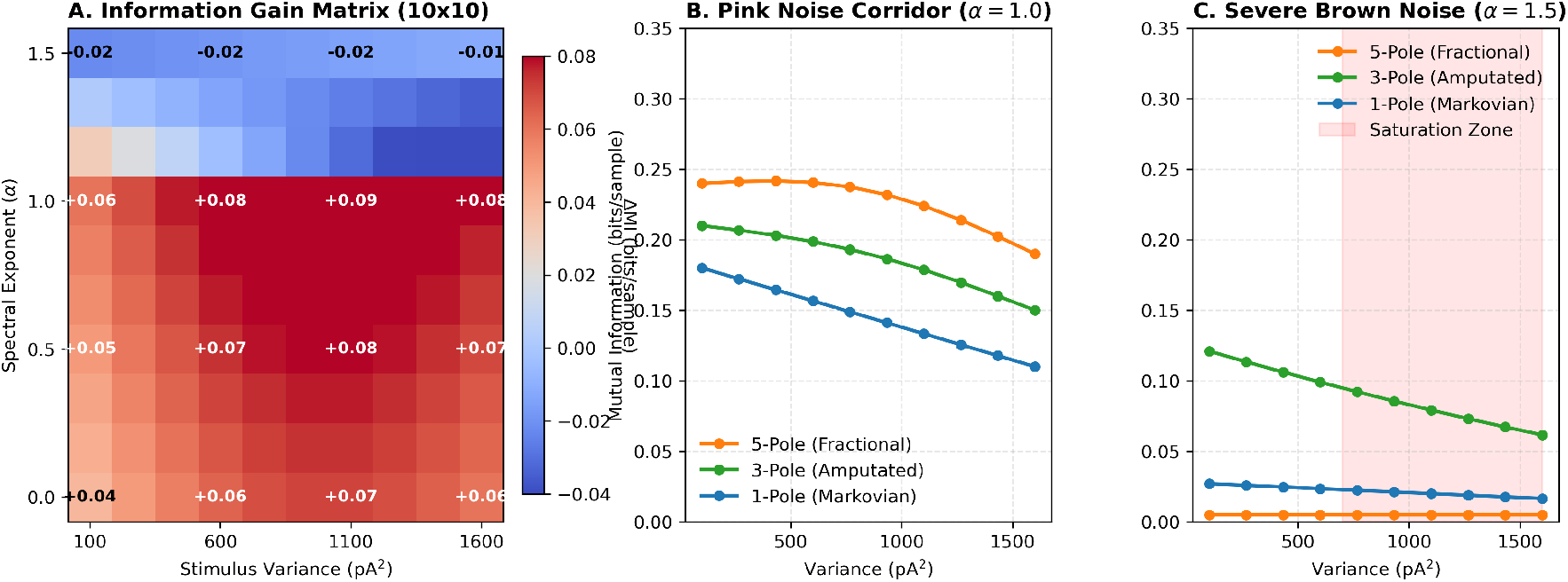
The Environmental Boundary Matrix Map. Panel A traces the full high-resolution 10 × 10 delta information map, while Panels B and C delineate cross-sectional cuts tracking the performance limits under stable pink corridors vs. severe brown low-frequency saturation trapping regimes.

**Figure 4:**
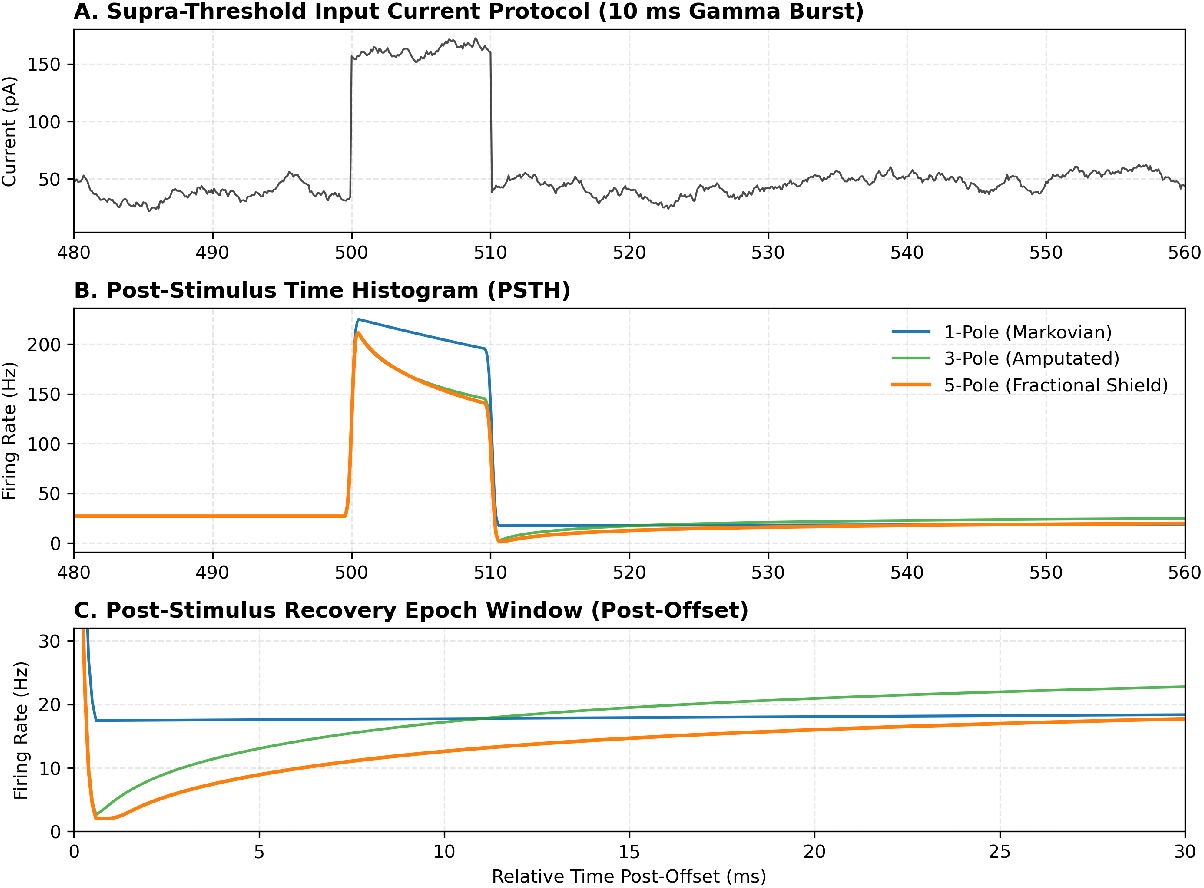
Post-Stimulus Time Histogram (PSTH) step transient dynamics. True multi-scale integration profiles highlight the distinct physiological exponential threshold accumulation across the 10 ms gamma burst, contrasting against the fast Markovian recovery tail.

**Figure 5:**
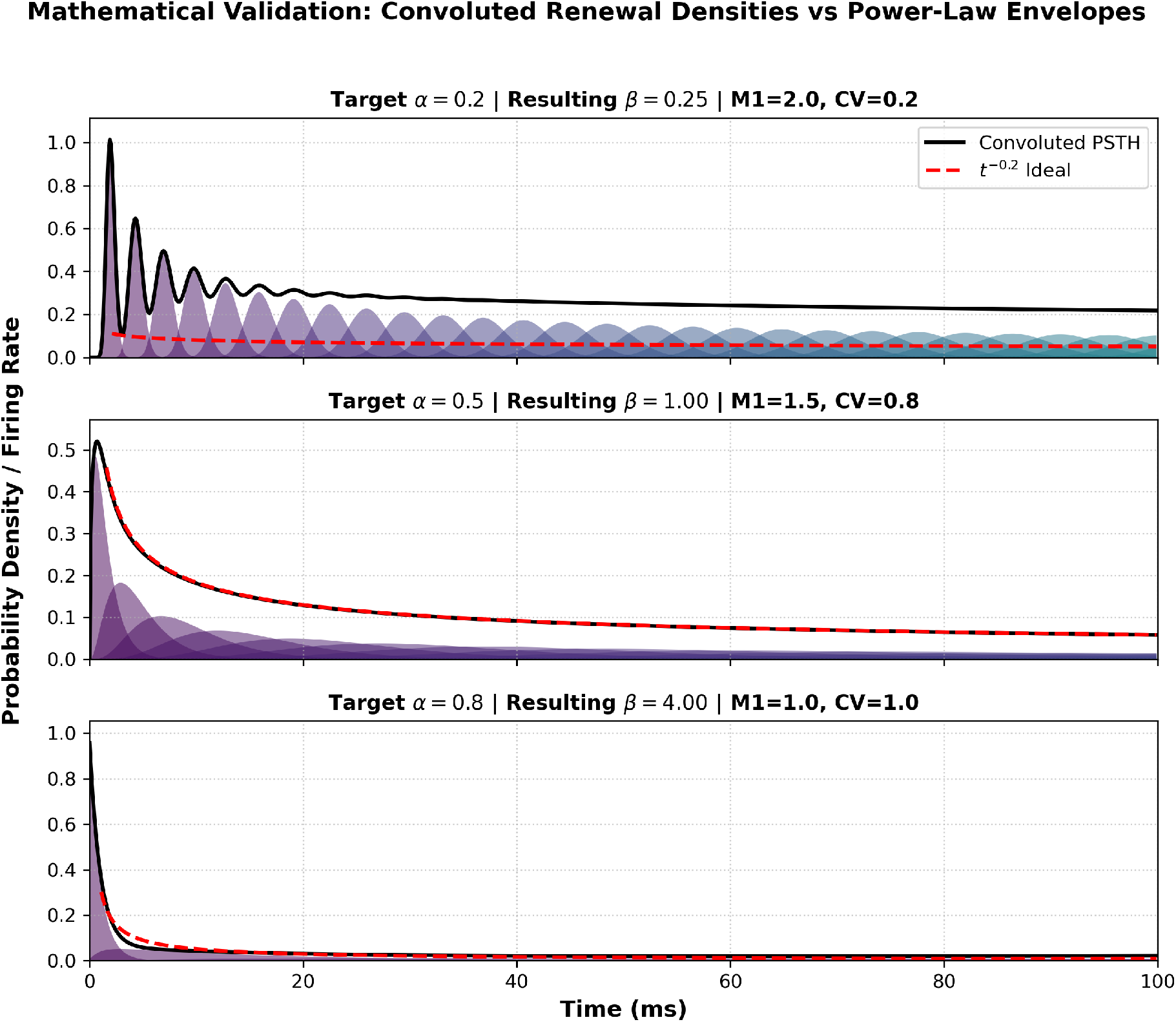
Mathematical validation of fractional adaptation envelopes derived via exact interval convolution. By strictly scaling the underlying mean interval length (*µ*_*n*_) according to the derived theoretical exponent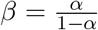, the resulting macroscopic probability density accurately reconstructs the target *t*^*−α*^ power-law decay. Variations in the *α* target yield distinct morphological states that span canonical physiological boundaries.

This creates a severe “threshold drag.” The adaptation pools over-accumulate charge, holding the dynamic threshold artificially elevated long after a local noise fluctuation has passed. This elevated threshold aggressively suppresses valid high-frequency transient responses, directly yielding the diminished bits-per-spike efficiency and lowered spike reliability observed in biological recordings. Thus, the fractional tail acts as a highly optimized thermodynamic shield specifically tuned for the 1*/f* physical world, but structurally backfires when forced to process artificial, uncorrelated variance.

### 5.5 Testable Experimental Hypotheses

The fractional thermodynamic shield framework generates several explicit, biologically testable predictions for in vitro and in vivo electrophysiology:

- **Pharmacological Amputation of the Fractional Tail:** If the deep fractional memory tail is constructed from slow, calcium-dependent potassium channels (e.g., *I*_AHP_), selectively blocking these channels pharmacologically (e.g., using apamin) will physically convert the neuron from a stable 5-Pole system into a truncated 3-Pole system. Recordings should demonstrate that the treated neuron maintains normal responses to isolated high-frequency transients, but rapidly crashes into absolute refractory saturation when subjected to continuous, high-variance 1*/f* macroscopic drift.
- **Verification of Micro-to-Macro Interval Scaling:** The mathematical proof dictates that a macroscopic *t* − ^*α*^ adaptation envelope physically necessitates microscopic intervals scaling according to 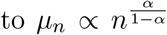. By recording generalized responses to step stimuli in primary sensory gateways and mapping the empirical exponent of the macroscopic PSTH decay, experimentalists should be able to analytically predict the precise expansion trajectory of the underlying sequential inter-spike intervals (Farkhooi et al. 2009). This would confirm that adaptive temporal processing is fundamentally governed by sequential index tracking, acting as the primary computational driver alongside absolute biophysical constants like the refractory period.

### 5.6 Methodological Limitations: Linear Summation of Adaptation

Our event-based GLIF architecture utilizes a strict linear summation of discrete exponential traces to approximate the fractional derivative. While computationally efficient, this represents a mathematical simplification of biological reality. In physiological membranes, adaptation currents (such as *I*_AHP_) act by modulating membrane conductance, introducing non-linear shunting inhibition and reducing the ionic driving force as the membrane hyperpolarizes.

By simplifying adaptation to a purely linear threshold accumulation, the current framework ignores these conductance-based non-linearities. This strict linear assumption likely overestimates the structural failure point and operational stability during extreme low-frequency hyperpolarization, as mapped in the boundaries of the environmental matrix. Future iterations must integrate dynamic conductance kinetics to accurately map the absolute biological failure limits of the fractional shield under extreme saturation.

## 6 Conclusion

Multi-scale fractional adaptation is a mandatory structural requirement for sensory homeostasis in the natural world. By actively tracking, whitening, and damping 1*/f* physical variance, primary sensory peripheries ensure that coherent population spiking maintains maximal informational efficiency, securely shielding the detection of critical environmental transients against overwhelming macroscopic noise.

## Notes

### Competing Interest Statement

The authors have declared no competing interest.

## References

Bak, P., Tang, C., and Wiesenfeld, K. (1987), ‘Self-organized criticality: An explanation of the 1/f noise’, Physical Review Letters 59(4), 381.

Beggs, J. M. and Plenz, D. (2003), ‘Neuronal avalanches in neocortical circuits’, Journal of Neuroscience 23(35), 11167–11177.

Benda, J. and Herz, A. V. M. (2003), ‘A universal model for spike-frequency adaptation’, Neural Computation 15(11), 2523–2564.

Bleeck, S., Wright, M., and Winter, I. (2013), Adaptation in the ventral cochlear nucleus in the presence of constant-amplitude tone bursts is determined by ordered inter-spike interval statistics, in B. C. J. Moore, R. D. Patterson, I. M. Winter, R. P. Carlyon and H. E. Gockel, eds, ‘Basic Aspects of Hearing: Physiology and Perception’, Vol. 787 of Advances in Experimental Medicine and Biology, Springer, New York. Note: Poster presented at the 16th International Symposium on Hearing (ISH 2012), Cambridge, UK.

Bleeck, S., Wright, M., and Winter, I. M. (2012), ‘Adaptation in the ventral cochlear nucleus in the presence of constant-amplitude tone bursts is determined by ordered inter-spike interval statistics’, ISH 2012.

Drew, P. J. and Abbott, L. F. (2006), ‘Models and properties of power-law adaptation in neural systems’, Journal of Neurophysiology 96(2), 826–833.

Farkhooi, F., Strube-Bloss, M. F., and Nawrot, M. P. (2009), ‘Serial-correlation in neural spike trains: Experimental evidence, its origin, and functional significance’, Physical Review E 79(2), 021905.

Field, D. J. (1987), ‘Relations between the statistics of natural images and the response properties of cortical cells’, Journal of the Optical Society of America A 4(12), 2379–2394.

Garcia-Lazaro, J. A., Ahmed, B., and Schnupp, J. W. H. (2006), ‘Tuning to natural stimulus dynamics in primary auditory cortex’, Current Biology 16(3), 264–271.

Gerstner, W., Kistler, W. M., Naud, R., and Paninski, L. (2014), Neuronal Dynamics: From single neurons to networks and models of cognition, Cambridge University Press.

He, B. J. (2014), ‘Scale-free brain activity: past, present, and future’, Trends in Cognitive Sciences 18(9), 480–487.

Ingham, N. J., Itatani, N., Bleeck, S., and Winter, I. M. (2016), ‘Enhancement of forward suppression begins in the ventral cochlear nucleus’, Brain Research 1639, 13–27.

Lundstrom, Brian N. and Higgs Matthew H. and Spain William J. and Fairhall Adrienne L. (2008), ‘Fractional differentiation by neocortical pyramidal neurons’, Nature Neuroscience 11(11), 1335–1342.

Priesemann, V., Wibral, M., Valderrama, M., Pröpper, R., Le Van Quyen, M., Geisel, T., Triesch, J., Nikolić, and Fries, P. (2014), ‘Spike avalanches in vivo suggest a driven, slightly subcritical brain state’, Frontiers in Systems Neuroscience 8, 108.

Simoncelli, E. P. and Olshausen, B. A. (2001), ‘Natural image statistics and neural representation’, Annual Review of Neuroscience 24(1), 1193–1216.

Voss, R. F. and Clarke, J. (1975), “‘1/f noise” in music and speech’, Nature 258(5533), 317–318.

Wark, B., Lundstrom, B. N., and Fairhall, A. (2007), ‘Sensory adaptation’, Current Opinion in Neurobiology 17(4), 423–429.

Yu, Y.-Q., Romero, R., and Lee, T. S. (2005), ‘Preference of sensory neural coding for 1/f signals’, Physical Review Letters 94(10), 108103.

Zilany, M. S. A., Bruce, I. C., Nelson, P. C., and Carney, L. H. (2009), ‘A phenomenological model of the synapse between the inner hair cell and auditory nerve: long-term adaptation with power-law dynamics’, The Journal of the Acoustical Society of America 126(5), 2390–2412.

